# Evolutionary and Structural Constraints Influencing Apolipoprotein A-I Amyloid Behaviour

**DOI:** 10.1101/2020.09.18.304337

**Authors:** RA Gisonno, T Masson, N Ramella, EE Barrera, V Romanowski, MA Tricerri

## Abstract

Apolipoprotein A-I (apoA-I) has a key function in the reverse cholesterol transport mediated by the high density lipoprotein (HDL) particles. However, aggregation of apoA-I single point mutants can lead to hereditary amyloid pathology. Although several studies have tackled the biophysical and structural impacts introduced by these mutations, there is little information addressing the relationship between the evolutionary and structural features that contribute to the amyloid behavior of apoA-I. We combined evolutionary studies, *in silico* mutagenesis and molecular dynamics (MD) simulations to provide a comprehensive analysis of the conservation and pathogenic role of the aggregation-prone regions (APRs) present in apoA-I. Sequence analysis demonstrated that among the four amyloidogenic regions described for human apoA-I, only two (APR1 and APR4) are evolutionary conserved across different species of Sarcopterygii. Moreover, stability analysis carried out with the FoldX engine showed that APR1 contributes to the marginal stability of apoA-I. Structural properties of the full-length apoA-I model suggest that aggregation is avoided by placing APRs into highly packed and rigid portions of its native fold. Following we set up to study the effect of natural mutations on protein conformation and stability. Compared to natural silent variants extracted from the gnomAD database, the thermodynamic and pathogenic impact of apoA-I amyloid mutations showed evidence of a higher destabilizing effect. MD simulations of the amyloid variant G26R evidenced the partial unfolding of the alpha-helix bundle with the concomitant exposure of APR1 to the solvent and the formation of beta-sheet segments at the C-terminus of apoA-I, giving a possible hint about the early steps involved in its aggregation. Our findings highlight APR1 as a relevant component for apoA-I structural integrity and emphasize a destabilizing effect of amyloid variants that leads to the exposure of this region. This information contributes to our understanding of how apoA-I, with its high degree of structural flexibility, maintains a delicate equilibrium between its monomeric native structure and intrinsic tendency to form amyloid aggregates. In addition, our stability measurements could be used as a proxy to interpret the structural impact of new mutations.

## Introduction

Apolipoprotein A-I (apoA-I) is the most abundant protein component of high-density lipoproteins (HDL) and is responsible for the reverse cholesterol transport from extracellular tissues back to the liver (Lund-Katz and Phillips 2010; Rader et al. 2009), which has been associated with a protective function against cardiac disease and atherosclerosis (Navab et al. 2009; Rosenson et al. 2015). The scaffolding functions of apoA-I in the HDL particle and its multiple protein-protein interactions, mainly with the lecithin:cholesterol acyltransferase and the ATP-binding cassette A1 transporter (Chroni et al. 2003; Manthei et al. 2020), forces it to maintain a dynamical and flexible conformation (Gursky and Atkinson 1996).

In contrast to these physiological functions, several point mutations affecting apoA-I have been associated with hereditary amyloid pathology (Sipe et al. 2016). These mutations are mainly distributed into two “hot spots,” located at the N-terminal region and the C-terminus of the protein, each one with a typical clinical manifestation (Das and Gursky 2015). Mutations that occur at the N-terminal region (residues 26–100) are characterized by amyloid deposits in the liver and kidney (Mucchiano et al. 2001; Obici et al. 2006), while those located at a short C-terminus domain (residues 170–178) are mainly described as inducing heart, larynx and skin deposits (Gaglione et al. 2018). In non-hereditary amyloidosis, full-length apoA-I is detected deposited in the intima of severe atherosclerotic plaques or as diffuse patches as senile forms of amyloid. This process has been associated with aging, but it has also been described in chronic pathologies such as Alzheimer’s disease and type 2 diabetes mellitus (Westermark et al. 1995).

Amyloid behavior of some apoA-I N-terminal fragments has been attributed to the presence of aggregation-prone regions (APRs) in its sequence and, specifically, to an APR located at the N-terminus (Obici et al. 2006). A recent study has proved that four regions of human apoA-I sequence are capable of forming cross-beta structures (Louros et al. 2015). It has been hypothesized that amyloidogenic mutations or post-translational modifications could promote aggregation through destabilization of the partially disorganized structure of apoA-I, described as a molten globular state, followed by the exposure of APRs. In this sense, most studies addressing the effect of amyloid variants have focused on the biophysical and physiological consequences of specific mutants. However, our understanding of the relationship between apoA-I sequence determinants and its aggregation process remains limited.

In this study, through an evolutionary analysis we characterized the conservation of aggregation-prone regions in a broad dataset of vertebrates apoA-I sequences. Using the recently described full-length consensus structure (Melchior et al. 2017), we examined the structural properties of apoA-I that contribute to minimize the exposure of its constituent APRs. *In silico* saturation mutagenesis analysis of apoA-I demonstrated that the evolutionary-conserved APR1, comprising residues 14-19, contributes to the thermodynamic stability of the N-terminus and revealed a common destabilizing effect for amyloid-associated variants. Using molecular dynamics simulations, we studied the conformational and dynamical impact of different amyloid variants on the structure of full-length apoA-I. Altogether, our results suggest that APR1 is an evolutionary and structural conserved component that contributes to the stability of apoA-I structure. Mutagenesis data emphasizes the destabilizing effect of amyloid variants, which in the case of the G26R natural variant is linked to the solvent exposure of APR1 and the formation of a beta-sheet element at the C-terminus of the protein. This information is relevant to understand how a marginally stable, but metabolically active protein manages to initiate the formation of an amyloid structure and develops a severe pathology.

## Results

### Molecular evolution of apoA-I aggregating regions within Sarcopterygii

Given that apoA-I has four previously characterized APRs (residues 14-19 for APR1, 53-58 for APR2, 67-72 for APR3 and 227-232 for APR4), we asked if these amyloid regions could be relevant to the protein functionality in spite of their known pathogenic role (Louros et al. 2015). To tackle this question, first of all we decided to explore the evolutionary conservation of these motifs within apoA-I sequences of sarcopterygian organisms. Our analysis was restricted to this group in order to cover a wide range of species across the evolutionary history of apoA-I but excluding groups with large divergence times that could confound the results (Yousaf, Raza, and Abbasi 2015). Sequences were collected from the Ensembl database and a multiple sequence alignment (MSA) was constructed in order to identify the APRs present in other species based on the previously reported sequences for humans. We employed the Tango software to predict the sequence-based aggregation propensity of each one of the APRs and also computed the sequence conservation from the MSA based on the Shannon entropy (H). Our results suggest that the APR1 and APR4 has retained their amyloid propensity in more than 60% and 40% of the sequences of the dataset, respectively. On the other hand, APR2 and APR3 presented a non aggregation behavior in virtually all the sequences (Figure 1A). Regarding the sequence conservation of aggregation-prone regions, our data showed that the sequence entropy of APRs residues was statistically higher than the average H value for apoA-I (*P* value = 0.018, Mann-Whitney U Test). This suggest that apoA-I APRs can retain their aggregating behavior despite a lower sequence conservation (Figure 1B). Interestingly, the H values for the APRs seemed to be different from each other, with APR1 having the higher sequence conservation (Supplementary figure 1).

**Figure 1:**
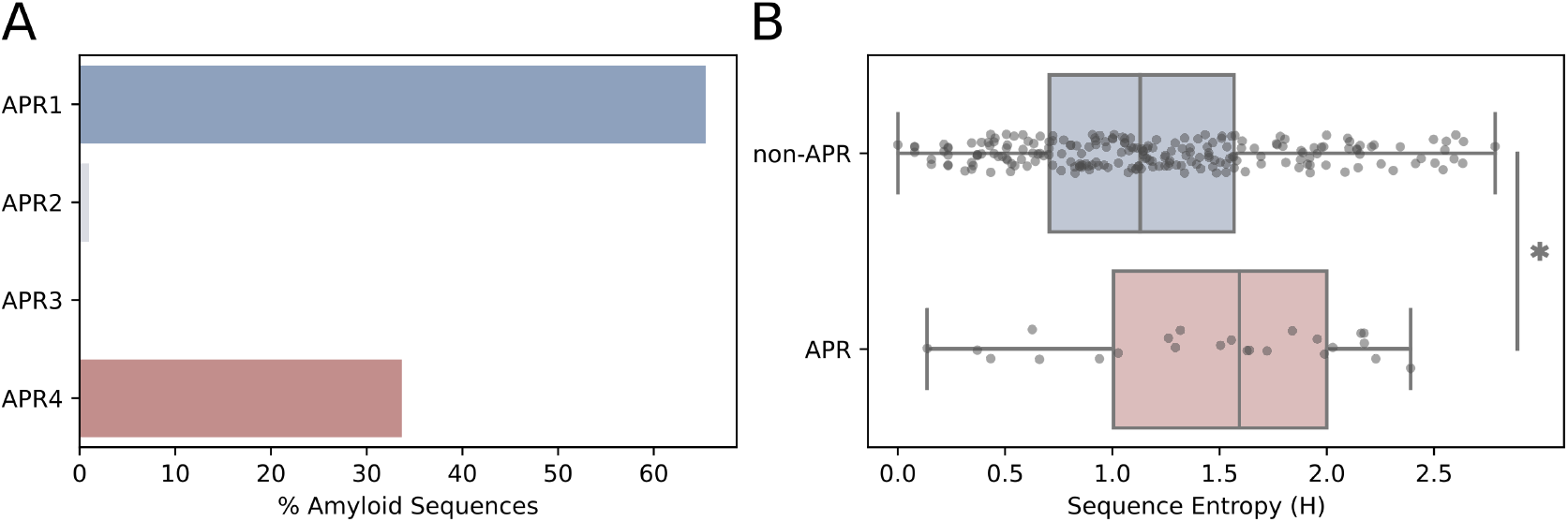
Evolutionary conservation of APRs within apoA-I sequences. **A** Percentage of sequences in our dataset that are described are amyloidogenic according to Tango (average score over 5%). **B** Sequence entropy (H) calculated for each residue inside the corresponding APR (P value = 0.018, Mann-Whitney U Test).

Motivated by the higher sequence diversity of APRs, we decided to investigate the selection constraints affecting apoA-I as a way to gain further insight into the conservation of its APRs motifs. A maximum likelihood phylogeny reflecting the evolutionary relationships between sequences was reconstructed from the MSA (Supplementary figure 2). Using this phylogeny as framework, we computed the site-wise evolutionary rates at the codon level (*dN/dS*, ratio of nonsynonymous to synonymous mutations) and evaluated its statistical significance in order to evidence the presence of selection constraints acting on the apoA-I sequence. In particular, we employed different methods from the HyPhy package in order to detect both pervasive and episodic selection events. In general terms, the evolutionary rate profile of apoA-I revealed that most of the protein sequence displayed dN/dS values between 0 and 1, indicating a predominance of negative and neutral selection regimes (Supplementary figure 3A). To depict the statistical evidence for the different types of selection constraints acting on apoA-I at residue-level, we used a cartoon representation for each of the HyPhy frameworks tested (FEL and FUBAR for pervasive selection, and MEME for episodic selection). In accordance with the entropy results, the residues corresponding to the APRs showed evidence of both purifying and neutral selection, implying that some APR residues tend to be conserved during evolution but others subject to sequence substitutions, either due a neutral or diversifying selection regime (Supplementary figure 3B).

### Structural modeling of apoA-I extant sequences

Prompted by the conserved amyloid behaviour of APR1 and APR4 albeit their not so strong sequence conservation, we decided to expand these results with information derived from protein structural data. We implemented an homology modeling approach to compare apoA-I structures corresponding to several extant sequences, including amphibians (*Xenopus tropicalis*), reptiles (*Crocodilus porosus* and *Chelonoidis abingdonii*), birds (*Gallus gallus*) and mammals (*Bos taurus, Canis lupus* and *Mus musculus*). To date, the most comprehensive and complete structure available for apoA-I corresponds to Dr. Davidson’s lab (ApoA-I consensus structure link), thus we used it as the template for our homology-based modeling pipeline. Modeller was used to generate a structural model for each target sequence and then we performed a step of energy minimization based on the FastRelax protocol from PyRosetta. An alignment of these modeled structures showed that the overall structure of apoA-I has been conserved among extant species. However, the alignment also displayed a significant conformational variability, mainly outside the helix bundle region (Figure 2A). Interestingly, all the modeled structures conserved similar profiles of intrinsic dynamics (represented by the mean squared fluctuation (MSF) of the alpha carbons of the protein backbone) and weighted contact number (WCN), a measure of how crowded is the molecular environment of a residue (Supplementary figure 4). This suggests that besides its conformational heterogeneity, apoA-I structures have conserved their overall intrinsic dynamics.

**Figure 2:**
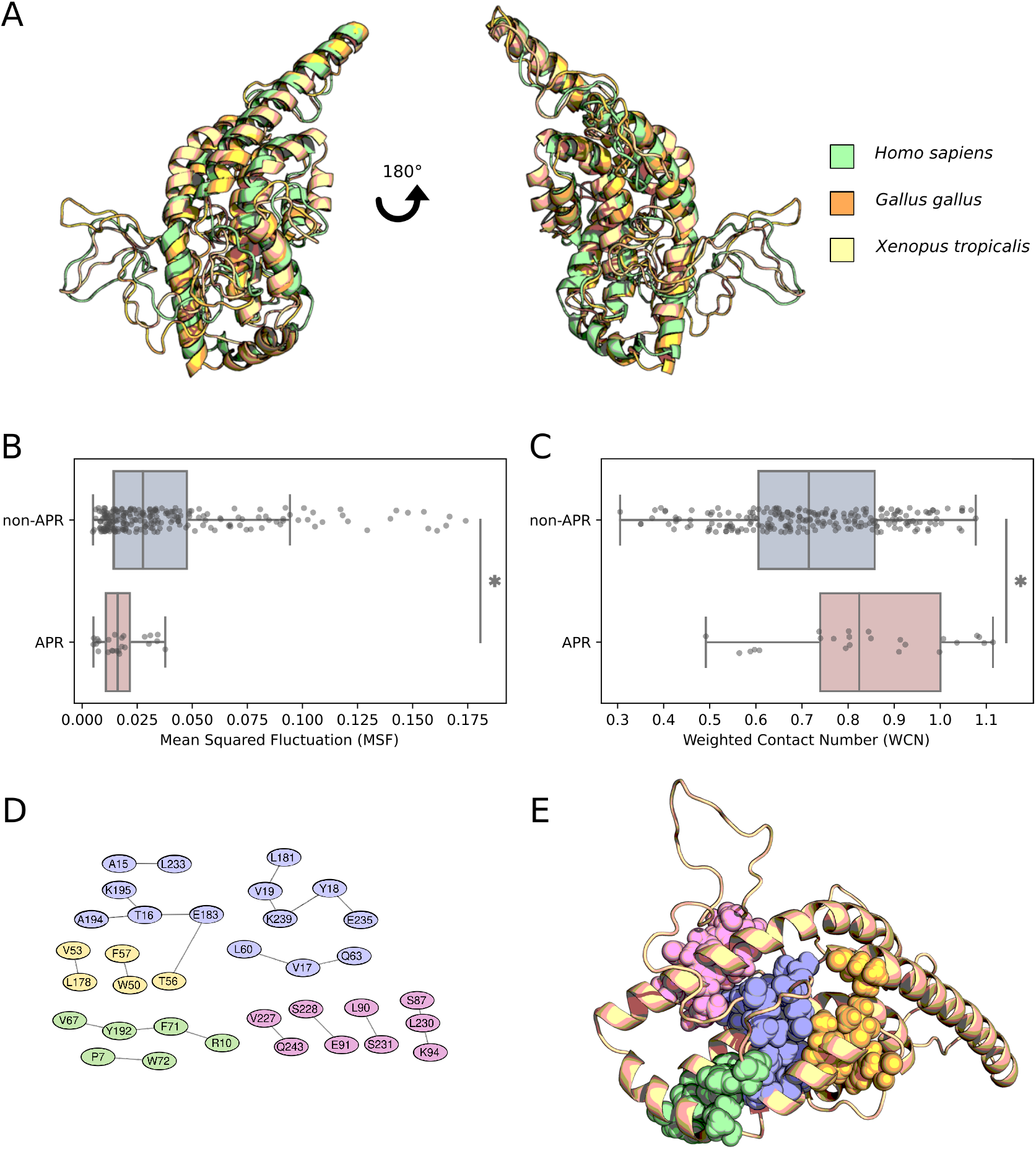
Dynamic and structural properties of apoA-I homologous structures. **A** Alignment of modeled structures corresponding to three extant apoA-I sequences. **B** Mean squared fluctuation (MSF; P value = 1.69 x 10^-4^) and **C** weighted contact number (WCN; P value = 1.23 x 10^-3^) corresponding to the APR and non-APR residues. **D** Network of residue contacts involving APRs, as computed by the Protein Contact Atlas and RING2 servers (APR1 in blue, APR2 in yellow, APR3 in green and APR4 in pink). **E** Structural mapping of apoA-I APRs contacts.

We used these structural models to further explore the intrinsic fluctuations levels and packaging numbers of the residue sites composing apoA-I APRs. Our results showed significantly lower MSF values for APRs residues when compared with the value distribution for the non-APR residues of apoA-I (Figure 2B, *P* value = 1.69 x 10^-4^, Mann-Whitney U Test). In a similar trend, the WCN values for APR residues were higher than non-APR regions on average (Figure 2C, P value = 1.25 x 10^-3^, Mann-Whitney U Test). In this structural context, APRs residues are integrated into relatively rigid and densely packaged portions of apoA-I, a hallmark of functionally relevant sites for the protein structure (Liu and Bahar 2012). To gain a deeper understanding about the molecular interactions that stabilized each APRs inside apoA-I structure, we used Protein Contact Atlas and RING2 servers to reconstruct the residue interaction network of each APR (Figure 2D and 2E). Based on this data, APR1 displayed the greater number of residues contacts, establishing interactions with helix H3 (residues 54-64) and 2 different regions of the C terminus (residues 183-195 and 235-239); this cluster of interactions are deeply buried inside the alpha-helix bundle of apoA-I. In contrast, the other APRs showed a smaller number of contacts, comprising more localized and solvent-exposed residues. Taking all these data together, although all APRs were characterized by low mobility and highly crowned molecular environment, APR1 seemed to be the greater contributor to the molecular interactions stabilizing the alpha-helix bundle region of apoA-I.

### Amyloid-associated variants have a destabilizing effect on apoA-I monomer structure

In order to better understand the contribution of APRs to apoA-I structural stability, we profiled the thermodynamic and pathological effect of every possible single point mutation in apoA-I sequence through *in silico* saturation mutagenesis. Destabilizing effect of each possible mutation in apoA-I sequence, represented by the difference in free energy (ΔΔG) between *wild type* and mutant structure, was measured using the FoldX empirical force field and the MutateX automation pipeline. To complement this approach, variant impact on protein function was estimated using Rhapsody. We noticed from the ΔΔGs distribution that most of the variants had a moderate impact on apoA-I stability (−1 kcal/mol < ΔΔG < 1 kcal/mol) (Figure 3A). Further examination revealed, as shown in Figure 3B, that apoA-I structure is highly sensitive to mutations in the region 7-28, which comprises APR1. (Complete FoldX results are available with Supplementary Figure 5). Rhapsody predictions also support this region as a mutation-sensible segment of apoA-I structure (Supplementary Figure 6). This result suggests that conservation of APR1 in apoA-I could be necessary to maintain the marginal thermodynamic stability of the alpha-helix bundle despite the risk to undergo aggregation. In line with our observations, APRs have been recently proposed to play a stabilizing role on protein structure (Langenberg et al. 2020).

**Figure 3:**
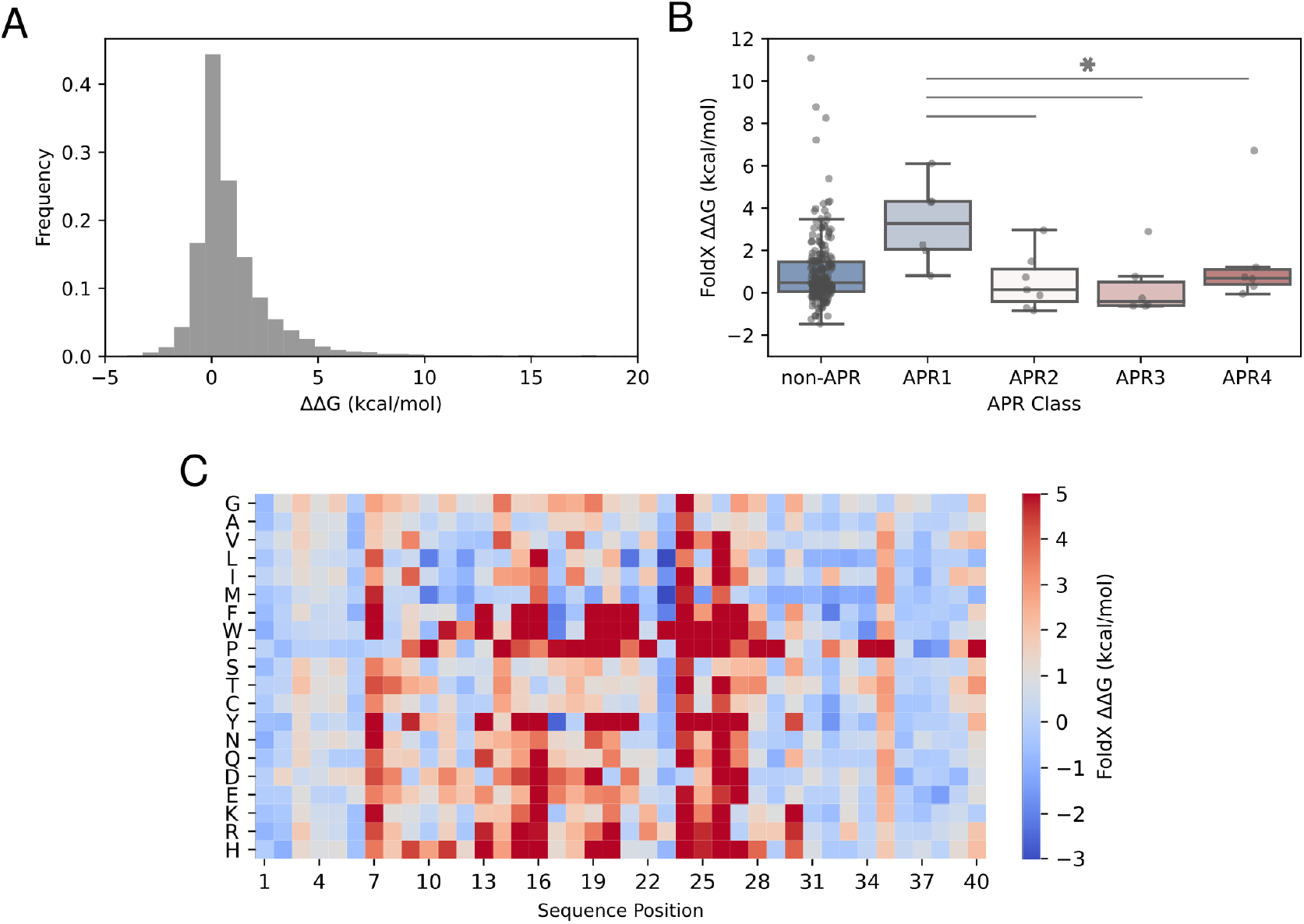
APR1 contributes to the stability of the alpha-helix bundle in apoA-I. The protein structural stability was quantified using the FoldX engine. The free energy difference (ΔΔG) was calculated by comparison between the ΔG of the mutant and *wild type* sequence A ΔΔG values distribution corresponding to all possible mutations. B Heatmap of ΔΔG values for the first 40 residues of apoA-I N-terminal region.

Given the observation that amyloidogenic variants do not modify the intrinsic aggregation tendency of APRs (Supplementary Table 1), as evidenced from TANGO predictions for the different apoA-I mutant sequences, we decided to investigate the impact of these variants on apoA-I stability. We used ΔΔG values to highlight differences between pathogenic variants associated with amyloid disease (Gogonea 2016) and natural variants reported by the gnomAD project (Karczewski et al. 2020). Our results evidenced that amyloid mutations had a destabilizing effect and a pathogenicity score significantly greater when compared with non-amyloid natural variants (Figure 4A and 4B), suggesting a close link between structural destabilization of apoA-I native conformation and the onset of amyloid pathology. As it was observed that a small group of variants in the gnomAD database showed an elevated impact on protein stability (> 2 ΔΔG kcal/mol), we decided to investigate how frequently they occur at population level. Figure 4C shows that variants with a severe impact on protein stability were present at low frequencies, thus minimizing their deleterious effect on the population. In contrast, variants with the higher frequency in our dataset had a nearly neutral effect on stability. It is worth noting that although gnomAD excluded subjects with mendelian and pediatric diseases from its cohorts, we cannot rule out the possibility that some of these destabilizing variants correspond to non diagnosed pathologies.

**Figure 4:**
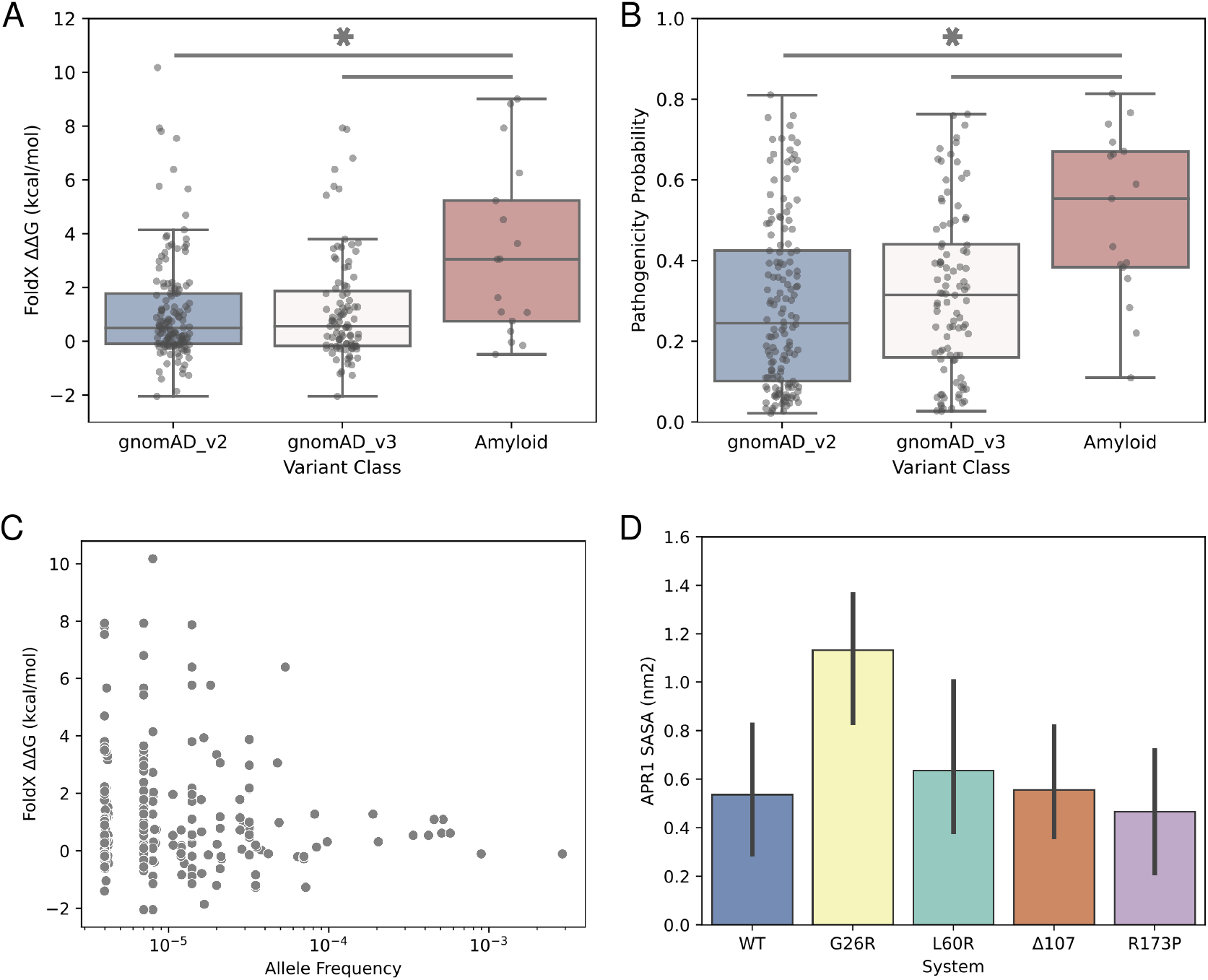
Impact of apoA-I variants on protein stability and function **A**-**B** Free energy difference (ΔΔG) and Rhapsody pathogenicity distributions for amyloid and gnomAD variant classes (P value < 0.01, Mann-Whitney U Test). **C** Allele frequency distribution for gnomAD variants as a function of their predicted effect on protein stability. **D** Molecular dynamics simulations of full-length apoA-I mutants. Solvent accessible surface area (SASA) calculated for the APR1 (residues 14-19). The SASA calculation for G26R displayed a higher APR1 exposure when compared with the *wild type* system (P value < 0.05 Student’s Test).

To complement our previous results showing the destabilizing effect of amyloid variants, we decided to study the dynamic properties of apoA-I amyloid mutants by conducting coarse-grain molecular dynamics simulations under the SIRAH force field. We selected four amyloid mutants (G26R, L60R, Δ107 and R173P) previously characterized by our group (Gaddi et al. 2020; Gisonno et al. 2020; Ramella et al. 2012; Rosú et al. 2015), plus the *wild type* protein, to prepare our simulation systems. Our selection also ensured that mutations were distributed throughout the apoA-I sequence. In the first place, we explored the overall dynamics of our systems by means of their root mean square deviation (RMSD). The recently described consensus structure for apoA-I was used as reference coordinates for RMSD calculations. We observed a great variability in the RMSD values for all simulated systems (5.4-10 angstrom) during our 1 microsecond simulation, which could be related with the highly dynamic and marginally stable structure proposed for apoA-I and is in agreement with the structural alignment of our modeled structures. We did not evidence any significant differences between the RMSD values of the different systems (Supplementary Table 2), suggesting that the impact of point mutations is negligible when compared against the intrinsic backbone dynamics.

Given the structural variability evidenced by RMSD, MD observables were computed over the last 100 ns of the simulations. Position-specific RMSF for each of the systems studied showed that loop regions 120-150 and 180-200 were the most flexible regions in apoA-I, while the N-terminal domain maintained a more compact structure during the simulation time (Supplementary Figure 7). These results are in good agreement with the MSF values computed with the GNM model, reinforcing the dynamic profile obtained for apoA-I. The similar fluctuation profiles between the *wild type* apoA-I and the abovementioned mutants suggest that mutations do not introduce major dynamical changes, at least during the simulation time frame. We explored the possible role of mutations in amyloid aggregation of full length apoA-I by analyzing the solvent accessible surface area (SASA) of each APR in our five systems. A significant increase in the solvent exposure of the APR1 was detected in the G26R system when compared against the *wild type* system, which suggests that some amyloid variants could lead to the solvent exposure of the aggregating regions (Figure 4D). Additionally, the formation of a beta-sheet structure was observed at the APR4 during the simulations of the system G26R (Supplementary figure 8). The low impact of the L60P, Δ107 and R173P variants on APRs exposure suggest that these mutants could affect other regions of apoA-I structure or may require further post-translational modifications in order to undergo amyloid aggregation.

## Discussion

Molecular mechanism of amyloid aggregation linked to apoA-I remains largely unknown, due in part to the limited structural information given its inherent conformational plasticity (Gursky and Atkinson 1996). This work builds upon evolutionary, dynamical and structural features of apoA-I in order to provide a comprehensive characterization of the amyloid phenomena in this protein, complementing the extensive experimental evidence available. Collectively, our results suggest an intimate relationship between aggregating regions and structural stability in apoA-I. Additionally, MD simulations of full-length apoA-I mutants shed light on the first steps of the aggregation process in some amyloid mutants.

The fact that apoA-I has conserved aggregating segments (APR1 and APR4) consistently along its evolutionary history raises questions about their structural relevance. Amyloid motifs have been proposed to contribute to protein structural stability through extensive interactions inside protein hydrophobic cores (Tartaglia and Vendruscolo 2010; Langenberg et al. 2020), which establish a trade-off between protein environment, foldability and aggregation propensity (Linding et al. 2004; Monsellier et al. 2008). Based on its conserved nature and FoldX stability results, it is possible to hypothesize that APR1 is necessary to ensure the marginal stability of apoA-I alpha-helix bundle, even though this region could trigger aggregation upon solvent exposure or proteolytic cleavage (Arciello, Piccoli, and Monti 2016). In addition, APR2 can act as a synergizing factor that aggravates the amyloid behaviour of apoA-I N-terminal peptide, albeit it low aggregation propensity (Wong et al. 2012; Mizuguchi et al. 2019). In this context, the structural features of APR1 (low intrinsic flexibility, highly packaged environment and several residue interactions) are likely to control its exposure to solvent and prevent aggregation events. Hydrogen-deuterium exchange experiments (Das et al. 2016) support the highly packaged nature of the alpha-helix bundle and the low solvent exposure of APR1 in apoA-I.

Amyloidogenic variants are primarily located towards the N-terminus of apoA-I, whereas variants associated with HDL deficiencies are clustered in the H5-H7 region (Gogonea 2016; Matsunaga et al. 2010). Through a comprehensive evaluation of the destabilizing effect and pathogenicity of each possible mutation affecting apoA-I we demonstrated that amyloid variants have a significant destabilizing effect on the monomer structure. The fact that TANGO aggregation tendency of APRs was not modified by the introduction of amyloid mutations, supports the hypothesis that aggregation propensity per se has a limited impact on the aggregation process of full-length apoA-I and certain destabilizing factors are required to *initiate* the amyloid process (Raimondi et al. 2011). In addition, we believe that ΔΔG values derived from our *in silico* saturation mutagenesis would be useful as a proxy for the initial study of novel apoA-I mutants. Taking advantage of the recently described consensus model of apoA-I (Melchior et al. 2017), our MD simulations of mutant G26R revealed a partial unfolding of the N-terminal alpha-helix bundle and a significant increase in the exposure of APR1, which is also congruent with the destabilizing effect predicted from our ΔΔG calculations. This partial unfolding is in line with the experimental reports of increased susceptibility to proteases (Adachi et al. 2012) and greater hydrogen-deuterium exchange rate of the alpha-helix bundle (Das et al. 2016) for this mutant. Moreover, beta-sheet secondary structures present at APR4 could provide a template for the aggregation of full-length apoA-I (Das et al. 2014).

Altogether, our results obtained from full-lenght protein support the current hypothesis that unfolding of the helix bundle and exposure of aggregating regions represents the first steps of apoA-I-mediated amyloidosis (Mizuguchi et al. 2019). The mild effect of L60R, Δ107 and R173P variants on apoA-I structure and APRs exposure suggest that further modifications could be required to promote protein aggregation of these mutants, like oxidation or proteolytic cleavage (Witkowski et al. 2018; Chan et al. 2015). Recently, the connection between the pro-inflammatory microenvironment and the formation of aggregation-prone species has been deeply characterized, reinforcing this hypothesis (Gisonno et al. 2020). Moreover, the presence of the N-terminal proteolytic fragment (residues 1-93) within patients’ lesions raises the hypothesis that mutations may facilitate the cleavage of apoA-I by circulating proteases (Cavigiolio and Jayaraman 2014; Kareinen et al. 2018). In agreement with the late onset of the hereditary apoA-I amyloidosis in patients, it may be hypothesized that mild chronic events may be required to induce the protein unfolding.

## Materials and Methods

### Evolutionary analysis of apoA-I sequences

A comprehensive dataset of sequences was generated by collating apoA-I orthologs available at Ensembl and Refseq databases (O Leary et al. 2015; Yates et al. 2019). To exclude low quality data, only sequences which did not contain ambiguous characters, had a proper methionine (M) starting codon and were longer than 200 amino acids were kept. Additionally, as both Ensembl and Refseq have overlapping data for some species, CD-HIT clustering tool (Fu et al. 2012) was employed to generate groups of similar sequences with an identity cut-off value of 0.98. Our final dataset comprised 104 protein sequences covering the Sarcopterygii lineage of Vertebrata.

In order to reconstruct a maximum likelihood phylogeny, a multiple sequence alignment (MSA) was built from the protein sequences using ClustalO with default parameters (Sievers et al. 2011) and the phylogenetic inference was carried out with the IQ-TREE software (Bui Quang Minh et al. 2020). The substitution model was selected based on the ModelFinder evolutionary model fitting tool (Kalyaanamoorthy et al. 2017) and the ultrafast bootstrap implemented in IQ-TREE was used to calculate the support values for phylogeny branches (B. Q. Minh, Nguyen, and Haeseler 2013). A supplementary phylogeny was reconstructed using a MSA generate with MAFFT in order to verify that the results obtained are independent of the aligning tool (Katoh 2002). Treefiles are available at the GitHub repository.

Visualization of the resulting phylogeny was carried out using the iTOL server (Letunic and Bork 2019).

### Selective pressure acting on apoA-I sequence

Nucleotide coding sequences were retrieved for each protein in our dataset using the NCBI Entrez eutils tools for Refseq sequences and the Ensembl orthologs dataset. Because the evolutionary rate estimation requires a codon-level alignment, the software PAL2NAL was used to align codons in nucleotide sequence using a protein alignment as a guide (Suyama, Torrents, and Bork 2006). The Hypothesis Testing using Phylogenies (HyPhy) package was used to conduct evolutionary analysis on the codon-based alignment. Before testing for evidence of selective pressure, we conducted a recombination analysis using the Genetic Algorithm Recombination Detection (GARD) method, (Pond et al. 2006) in order to screen for possible recombination events in our alignment; it is known that the presence of recombination leads to a larger number of false positives in selection analysis. We inferred the natural selection strength (Omega, *dN/dS*) for each alignment position using our phylogeny as framework. We employed the Fixed Effects Likelihood (FEL) (Pond and Frost 2005) and the Fast Unconstrained Bayesian Approximation (FUBAR) methods (Murrell et al. 2013) to quantify the dN/dS ratio for each codon in the alignment. Although both methods provide similar information, FEL provides support for negative selection (dN/dS < 1) whereas FUBAR has more statistical power to detect positive selection (dN/dS >1). Because codon alignment positions are difficult to put in structural context, data were extracted for codons occurring in *wild type* human apoA-I.

### Comparative structural modeling of apoA-I extant sequences

Structural models of apoA-I based on extant sequences were obtained with the Modeller software (Šali and Blundell 1993). A MSA between the target and template sequences was used to guide the modeling process and the consensus structure of the human apoA-I was selected as reference (Melchior et al. 2017). The raw models obtained with Modeller were subjected to a step of energy minimization using FastRelax from the PyRosetta suite in order to sample low-energy conformations that could potentially resemble the native state of the protein (Chaudhury, Lyskov, and Gray 2010). The resulting structures were visualized and aligned with the PyMol Molecular Graphics System, Version 2.0 Schrödinger, LLC.

Packaging level for residue *i* was represented by its Weighted Contact Number (WCN), which was calculated as follows:

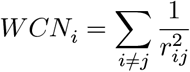

Where, *r_ij_* is the distance between the geometric center of the side-chain atoms for residue *i* and residue *j*. Calculations were carried out using a custom script developed by Sydykova et al. (Sydykova et al. 2018).

Protein intrinsic dynamics was characterized using a coarse-grained simulation model based solely on protein topological information represented as a Gaussian Network Model (GNM). In this approach, protein structure is modeled as a network of nodes (alpha carbons) connected by springs. Numerical resolution of this model allows the calculation of the equilibrium displacement for all nodes (Mean Square Fluctuation, MSF), describing the global motions of the system. The ProDy package (Bakan, Meireles, and Bahar 2011) was used to adjust a GNM to the apoA-I consensus structure. We selected the first ten slow modes for analysis and plotting, since they have been reported previously as the main determinants of the global dynamics of protein structure (Kitao and Go 1999). Residue interactions present within apoA-I structure were computed using two different approaches, RING2 and Protein Contact Atlas (Piovesan, Minervini, and Tosatto 2016; Kayikci et al. 2018). Briefly, both methods compute all non-covalent atom contacts between residues and use them to create a residue contact network were each node represents a residue of the protein and the edges between residues indicate the presence of at least one atomic contact.

### Conservation of Aggregation Prone Regions (APRs)

Signal peptide sequences were trimmed and removed from the MSA to retain only the mature protein sequence. TANGO software (Fernandez-Escamilla et al. 2004) was used to detect APRs in the protein sequences dataset. This algorithm predicts beta-aggregation using a space phase where the unfolded protein can adopt one of five states: random coil, alpha-helix, beta-turn, alpha-helical aggregation or beta-sheet aggregation. Importantly, TANGO is based on the assumption that the core regions of an aggregate are fully buried. Predictions were carried out using default settings: no protection for the C-terminus and N-terminus, pH 7, temperature of 310° K and ionic strength of 0.1. Output files provide an aggregation score per position; as suggested in the TANGO manual and elsewhere, contiguous regions comprising five or more residues with a score of at least five were annotated as an APR. To address the impact of single point mutations in apoA-I aggregation tendency we ran TANGO for each mutant sequence and compared the scores profile against the *wild type* sequence. TANGO software was downloaded from http://tango.crg.es using an academic license.

### Thermodynamics impact of missense variants

The FoldX engine (Guerois, Nielsen, and Serrano 2002) implements an empirical energy function based on terms significant for protein structure stability. The free energy of unfolding (ΔG) of the protein includes terms for van der Waals interactions, solvation of apolar and polar residues, intra and intermolecular hydrogen bonds, water bridges, electrostatic interactions and entropic cost for fixed backbone and side chains. Changes in free energy of folding upon mutation is calculated as the difference between the folding energy (ΔΔG) estimated for the mutants and the *wild type* variants. Although FoldX seems to be more accurate for the prediction of destabilizing mutations and less accurate for the prediction of stabilizing mutations, in both cases it was shown that FoldX is a valuable tool to infer putative relevant sites for structural stability. FoldX 5 suite was downloaded from http://foldxsuite.crg.eu/academic-license-info.

We employed MutateX software (Tiberti et al. 2019) to automate the prediction of ΔΔGs associated with the systematic mutation of each available residue within apoA-I, by employing the FoldX energy function. At the heart of MutateX lies an automated pipeline engine that handles input preparation and performs parallel runs with FoldX. Basic steps involve protein data bank (PDB) structure repair (involving energy minimization to remove unfavorable interactions), model building for the mutant variants, energy calculations for both mutant and *wild type* structures and summarizing the estimated average free energy differences.

### Pathogenicity scoring of missense variants

The Rhapsody prediction tool (Ponzoni et al. 2020) consists of a random forest classifier that combines sequence, structure, and dynamics-based features associated with a given amino acid variant and is trained over a comprehensive dataset of annotated human missense variants. Dynamical features include: mean-square fluctuations of the residue at the mutation site, which estimates local conformational flexibility; perturbation-response scanning effectiveness/sensitivity, accounting for potential allosteric responses involving the mutation site, and the mechanical stiffness at the sequence position of the mutated residue. These properties are computed from Elastic Network Models (ENM) representations of protein structures that describe inter-residue contact topology in a compact and computationally-efficient format that lends itself to a unique analytical solution for each structure. The algorithm was recently upgraded to include coevolutionary features calculated on conserved Pfam domains, and the training dataset was further expanded and refined. The latter combines annotated human variants from several publicly available datasets (Humvar, ExoVar, predictSNP, VariBench, SwissVar, Uniprot’s Humsavar and ClinVar). All analyses were performed using the Rhapsody server http://rhapsody.csb.pitt.edu/

### Molecular Dynamics Simulations

Coarse grained Molecular Dynamics simulations were performed with the SIRAH force field (Machado et al. 2019) and GROMACS 2018.4 software package (Abraham et al. 2015). We employed the consensus model of human apoA-I in its monomeric and lipid-free state, proposed by Davidson et al. (Melchior et al. 2017). The PDB file was downloaded from Davidson Lab homepage (http://homepages.uc.edu/~davidswm/structures.html). Mapping atomic to coarse-grained representations was done with a Perl script included in SIRAH Tools (Machado and Pantano 2016). G26R, L60R, R173P and Δ107 mutants were generated with Chimera (Pettersen et al. 2004), editing the coordinates of the consensus model pdb file. For the case of the deletion mutant, we removed Lys107 and connected residues Lys106 and Trp108 with an unstructured segment using Modloop (Fiser and Sali 2003). *Wild type* apoA-I and the mutant systems were assembled using the following setup: The protein was placed inside an octahedron simulation box defined by setting a distance of 1.5 nm between the solute and the edges of the box. Systems were solvated setting a 150 mM NaCl concentration following the protocol proposed by Machado et. al. (Machado and Pantano 2020). Energy minimization and heating steps were done following the protocol recommended by Machado et al. (Machado et al. 2019) using positional restraints in the protein backbone to ensure side-chain relaxation, especially in the mutant models. Production runs were performed by quintuplicate in the absence of any positional restraint, generating 1 microsecond trajectories at 310 K using a 1 bar NPT ensemble. Structural analysis was performed with GROMACS tools gmx rmsf, gmx gyrate and gmx sasa. Root mean square fluctuation was calculated for each residue aligning the full trajectory APOA-1 coordinates with the initial models. Radius of gyration and Solvent accessible surface areas (SASA) were obtained averaging the values corresponding to the last 0.1 microsecond of simulation. The SASA calculations were measured over three amyloid prone regions, comprising residues 14-19 (APR1), 53-58 (APR2), 67-72 (APR3) and 227-232 (APR4).

## Code and Files Availability

All Python packages used were installed through the Conda environment manager into a single environment. A requirements file is available in the repository of this project in order to install dependencies used in our analysis. The workflow manager Snakemake was used in the evolutionary analysis in order to gain reproducibility and consistency of the results (Koster and Rahmann 2012). The data, Snakefile and Python scripts used in this work are available at https://github.com/tomasMasson/APOA1_evolution.

## Statistical Analyses and Visualizations

Scipy Python library was used for data manipulation and all statistical analyses (Virtanen et al. 2020). Statistical significance was determined using Mann-Whitney U Test for variant’s impact comparison and Student’s Test for MD observables. MD graphs are reported as means ± standard deviation derived from five independent experiments. All visualizations were prepared with the Seaborn library (Waskom et al. 2020).

