## Supplementary Material for "Evolutionary and Structural Constraints Influencing Apolipoprotein A-I Amyloid Behaviour"

### Supplementary figure 1

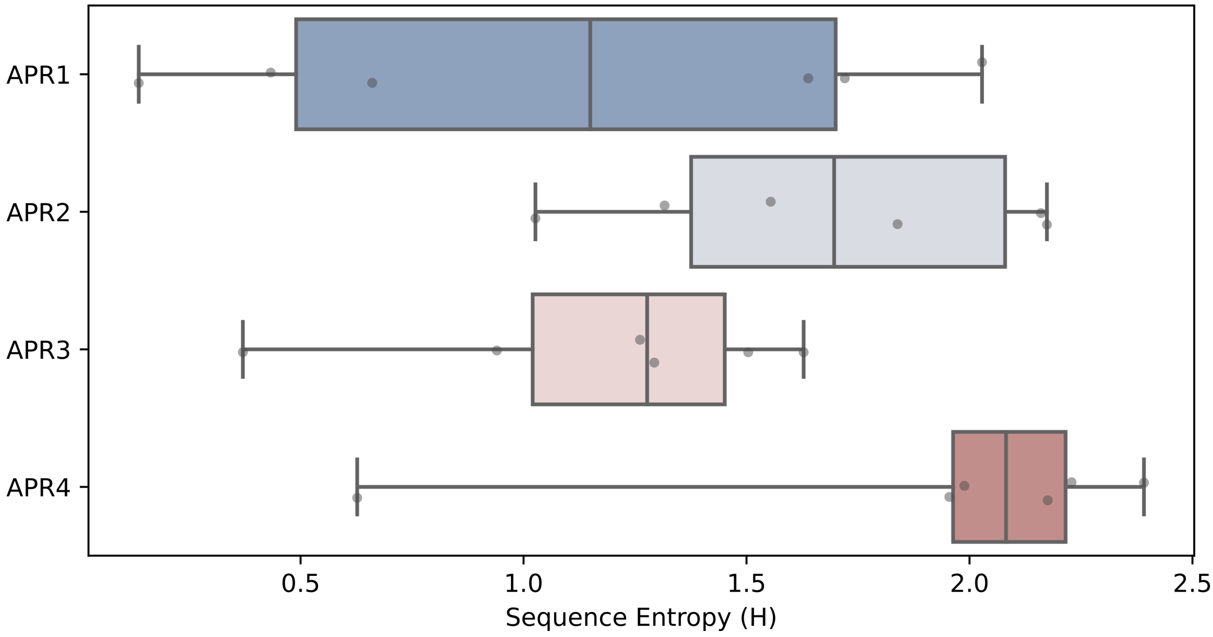


Figure 1: APRs Shannon's entropy (H) content.

### Supplementary figure 2

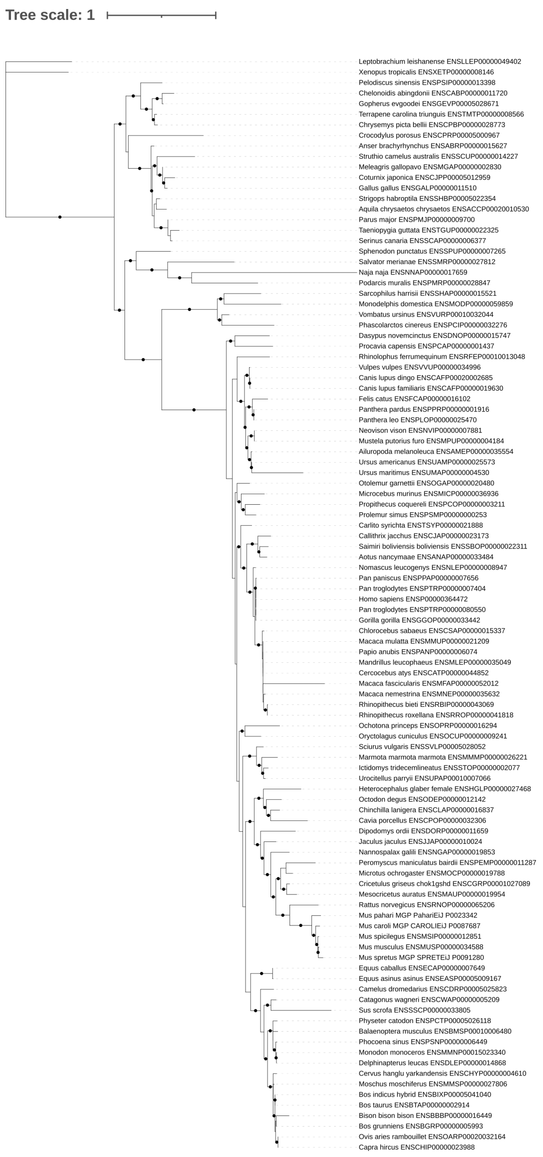


Figure 2: Maximum likelihood phylogeny inferred from a multiple sequence alignment of sarcopteriigyan apoA-I sequences. Black dots indicate node support values greater than 90% (calculated using 1000 replicates of UltraFast Bootstrap).

#### Supplementary figure 3

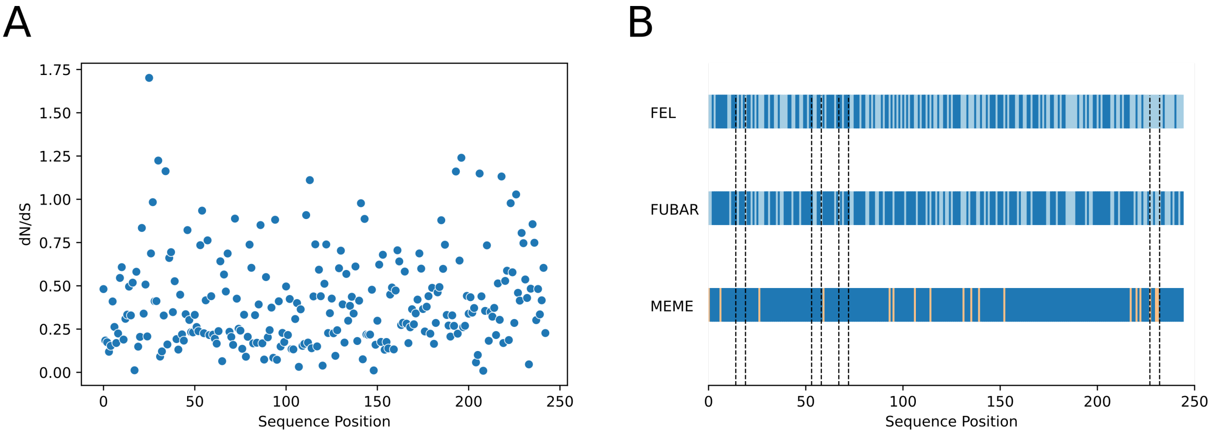


Figure 3: **A** Evolutionary rate (dN/dS) for each residue in the apoA-I codon alignment. **B** Cartoon depicting the evidence of adaptive evolution for each site of apoA-I sequence; residues under purifying, neutral and diversifying selection are colored in blue, paleblue and orange, respectively.

### Supplementary figure 4

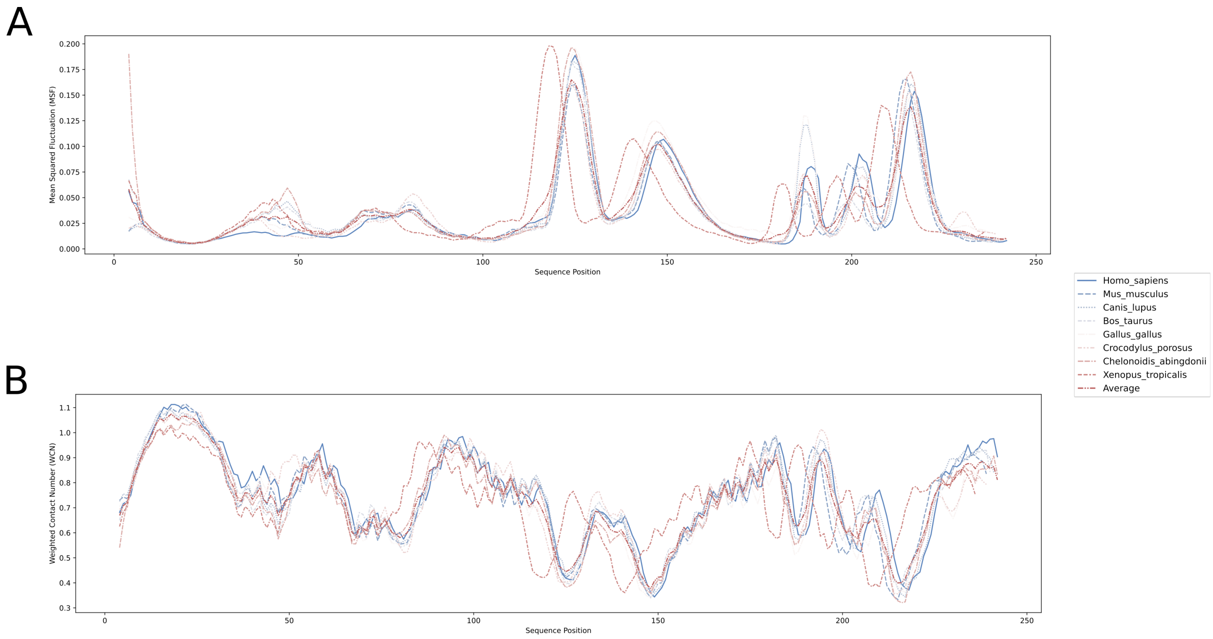


Figure 4: **A** Mean squared fluctuation (MSF) and **B** weighted contact number (WCN) profiles computed for each modelled structure

### Supplementary figure 5

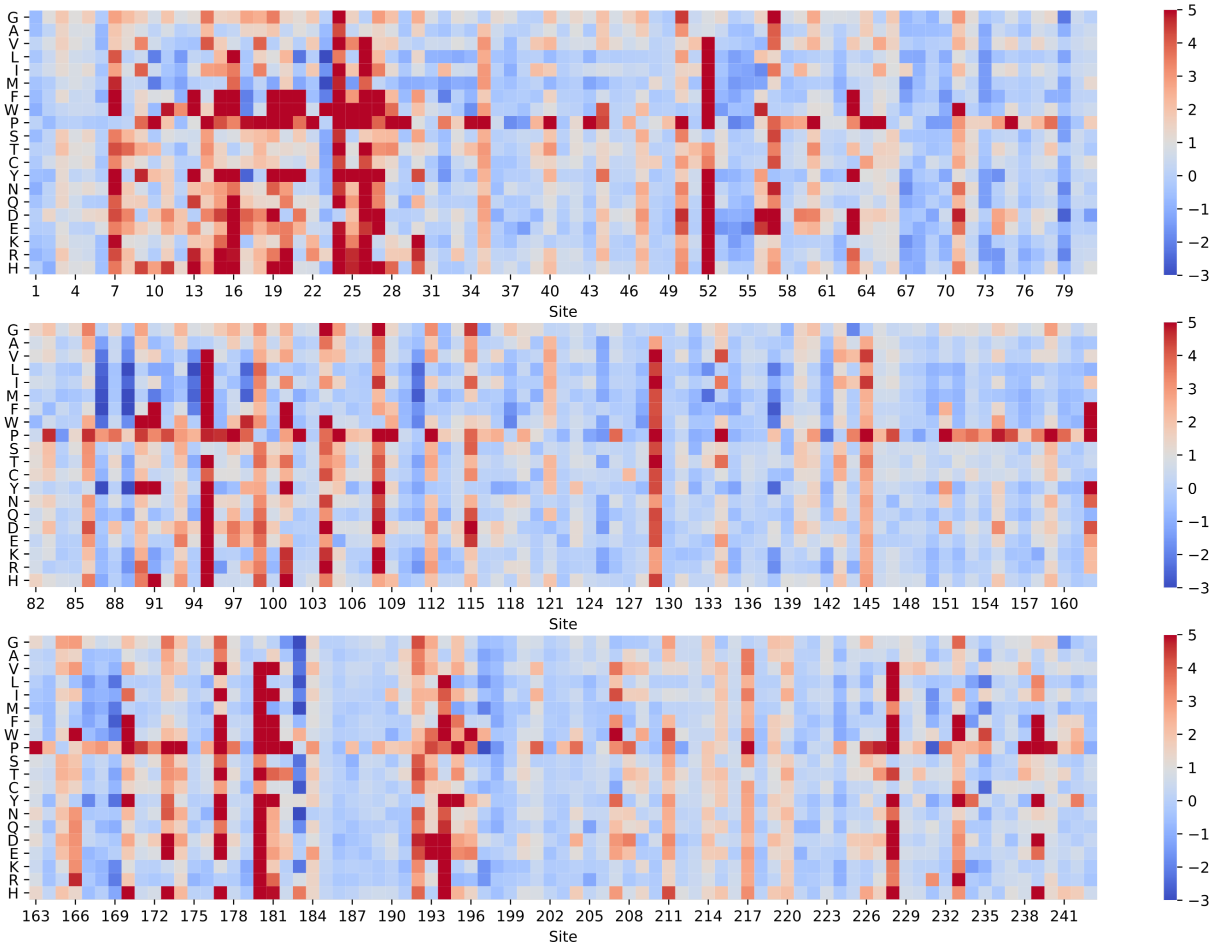


Figure 5: FoldX thermodynamic destabilization landscape. ΔΔG values obtained by *in silico* saturation mutagenesis of apoA-I structure using the FoldX engine. Mutation introduced is depicted in the Y axis. Scales at the right indicate ΔΔG values expressed in kcal/mol.

### Supplementary figure 6

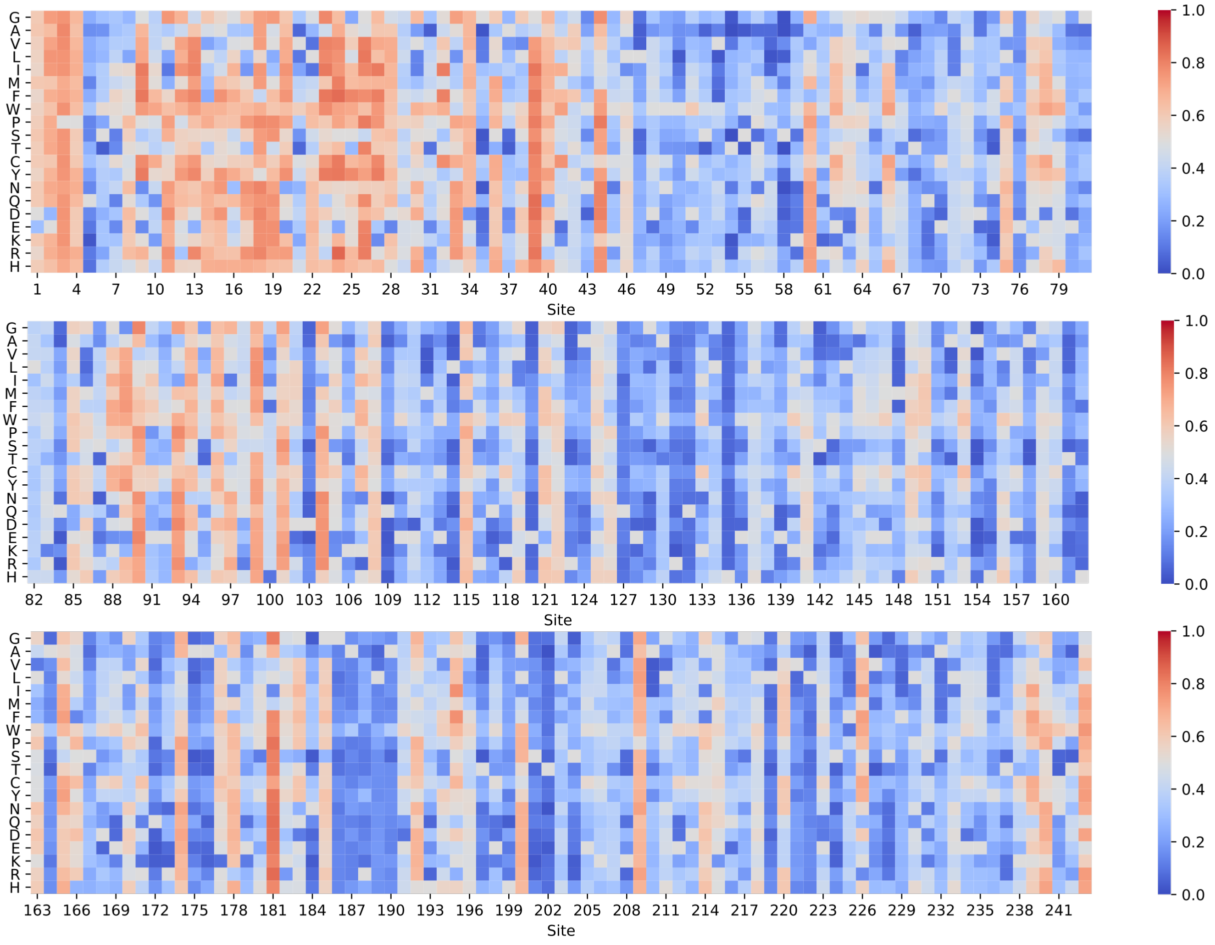


Figure 6: Rhapsody pathogenicity landscape. Pathogenicity values obtained by *in silico* saturation mutagenesis of apoA-I structure using the Rhapsody package. Mutation introduced is depicted in the Y axis. Scales at the right indicate pathogenicity score (1 more pathogenic, 0 less pathogenic)

### Supplementary figure 7

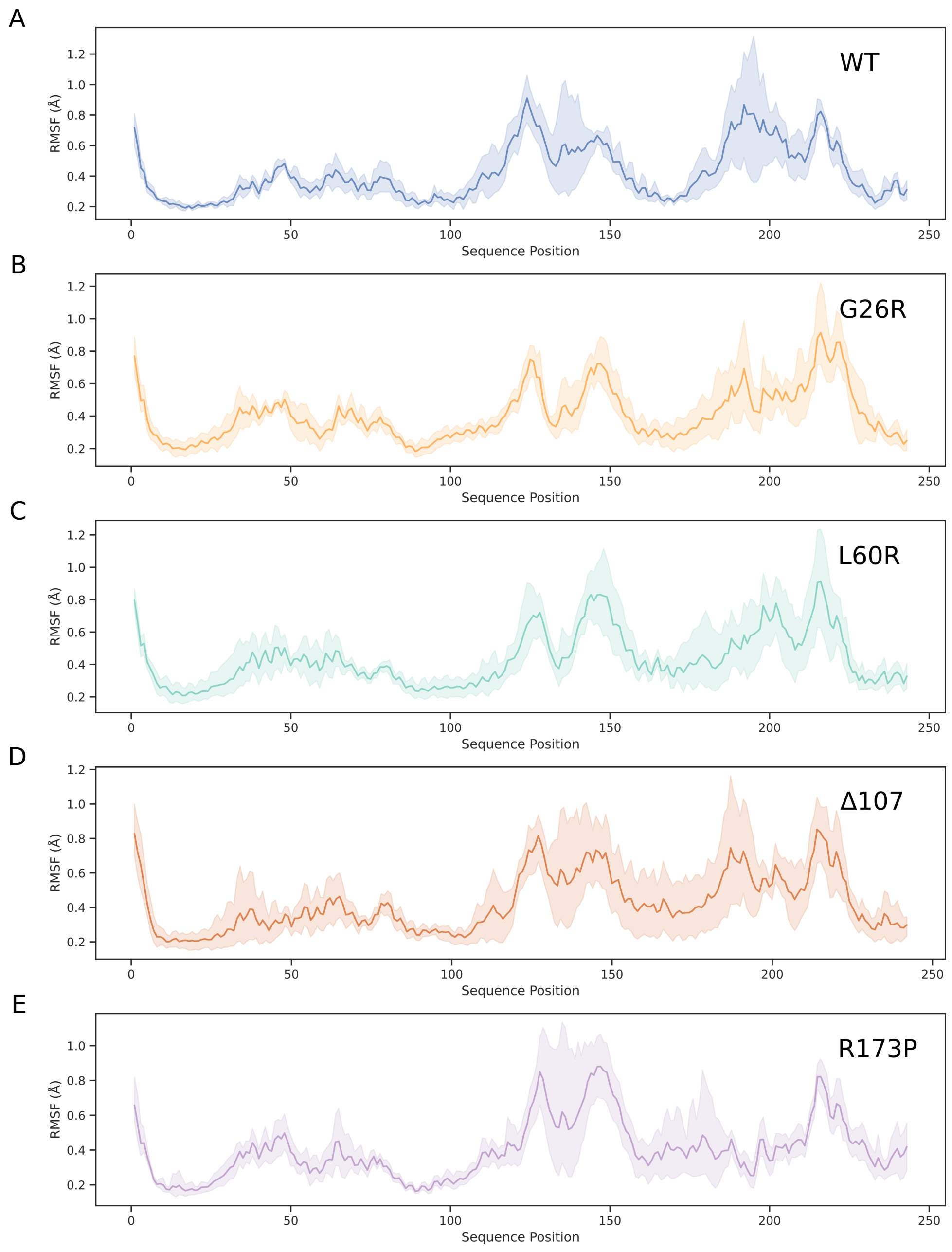


Figure 7: **A-E** Root mean square fluctuation (RMSF) profiles for apoA-I mutants. RMSF values were computed for each protein position over the last 100 ns of the simulation. Mean values are depicted together with its standard deviation.

#### Supplementary figure 8

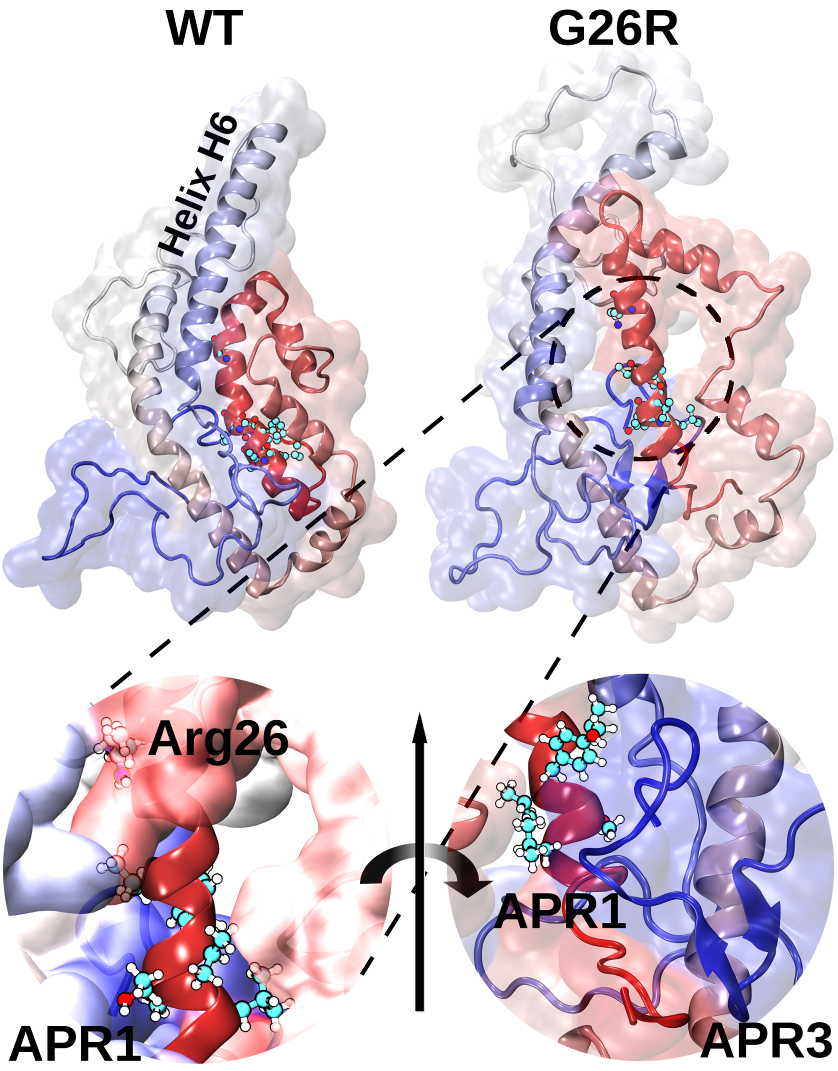


Figure 8: Graphical representation of the consensus model (WT) and the final snapshot of one of the replicas simulated for the G26R mutant. The substitution of glycine by arginine in position 26 destabilizes the helix bundle, expelling helix H6 with the concomitant solvent exposure of APR1 (left inset). A 180º view rotation shows the beta-sheet hairpin formed between residues S224 and A232, corresponding to the APR4 (right inset).
